# Micro-invasive probes for monitoring electrical and chemical neural activity in nonhuman primates

**DOI:** 10.1101/2025.01.30.635139

**Authors:** Usamma Amjad, Shreya Mahajan, Jiwon Choi, Ritesh Shrivastav, Raymond Murray, Abby Somich, Olivia Coyne, Helen N. Schwerdt

## Abstract

We leveraged carbon fiber materials for creating sensors that provide dual neurochemical and electrical neural activity recording at micro-invasive (10 µm) spatial footprints proximal to recording sites, and enabling these measurements from deep brain targets of primates with conventional cranial chambers. These shaft-assisted micro-invasive probes (s-µIPs) are approximately 10 µm in diameter along the distal length (1 – 15 mm) immediately surrounding the targeted recording site. This micro-invasive portion ensures that the recording site is isolated from tissue damage induced by the wider shaft portion of the device. The shaft (150 – 165 µm in diameter) within the device stiffens the remaining length of the probe (> 100 mm), and provides compatibility with standard intracranial insertion protocols (e.g., guide tubes and chamber setups) that require a sufficiently rigid and long shaft for deep brain insertion in monkeys. The s-µIP was further expanded to provide dual-channel chemical and electrical neural activity recording with micrometer spatial resolution. Measurements of reward- and movement-related dopamine, spikes, and local field potentials were made from single and dual-channel s-µIPs implanted in task-performing monkeys. Recordings from chronically implanted s-µIPs display the capability of functional multi-modal (chemical and electrical) neural activity measurements over 1-year post-implantation from micro-invasive devices.

## Introduction

Chronic recording of dopamine and other extrasynaptic neurotransmitters and neuromodulators requires sensors that minimize physical perturbation of the targeted tissue recording environment (*1*–*3*). Implanted probes induce a host of inflammatory responses including scar formation that physically block the diffusion of targeted neurochemicals to the sensor surface. These responses arise from the insertion process, as well as the physical indwelling of the sensor, which can further lead to motion induced trauma due to the mechanical mismatch between the sensor and the neural tissue. The insertion and prolonged indwelling of these sensors also disrupt the natural organization of cellular structures in the targeted tissue, potentially hindering the ability to accurately capture normal physiological chemical fluctuations. Thus, neurochemical recording sensors should be as small as possible to avoid physical displacement relative to the targeted recording site during movements and flexible to mechanically conform to the brain tissue (*4*). On the other hand, smaller sensors are more difficult to introduce into the brain, as they are usually in the form of a cylinder where bending stiffness is inversely proportional to its diameter (*EI* ∝ *D*^4^, where *E* is the Young’s Modulus, *I* is the area moment of inertia, and *D* is the diameter). Furthermore, flexibility scales with the length of the probe (*δ* ∝ *L*^3^, where δ is the deflection, and L is the length), making this even more difficult in animal species with larger brains (e.g., monkey or human). Here, we describe the process for building shaft-assisted micro-invasive probes (s-µIPs) that minimize trauma near the recording site, while incorporating a stiff shaft to facilitate insertion towards deep brain targets for application in primates.

Micro-invasive probes (µIPs) have been developed to help address these issues and have enabled year-long and longitudinally stable measurements of dopamine in rodents (*2*). The µIPs have a maximal diameter of 8 –10 µm, which is on the same size scale as the average neuron cell body (< 10 µm). Furthermore, these sensors have been shown to produce little to no visible inflammatory response when inserted in the rodent brain (*2*). The µIPs are made of carbon fiber (CF) (5 – 7 µm diameter), which acts as the electrochemical interface for measuring current generated in response to reduction and oxidation (i.e., redox) of targeted molecules.

Standard CF electrodes (CFEs) are typically made by threading the CF into a larger (∼ 90 – 100 µm diameter) glass or silica tube for insulation, leaving just the tip exposed for neural recording. Such CFEs have been used successfully in many seminal studies investigating dopamine’s role in motivation, learning, and other critical behaviors in rodents (*5, 6*), monkeys (*7*), and humans (*8*). The goal of the µIP design is to minimize the spatial footprint and resulting flexural rigidity (*K* < 8 × 10^−11^ N m^2^ as compared to the 90 µm silica tube with a *K* > 2.3 × 10^−7^ N m^2^) by leveraging the inherently small CF and the application of a thin (0.5 – 1.3 µm) conformal polymer insulation (i.e., parylene-C, py). This smaller sized and more flexible probe reduces the trauma induced by its implantation. The thinner insulation also facilitates increased channel capacity, since more probes can be introduced without significant accrual of induced trauma. On the other hand, the py-coated CF is highly flexible and difficult to insert into the brain on its own. In rodents, biodissolvable coatings are used to temporarily stiffen such probes to enable brain insertion (*9*). These biodissolvable coatings may not be directly compatible with standard neurophysiological setups used in monkeys, where chambers are typically installed to provide a window for accessing the brain with electrodes and guide tubes are used to penetrate the stiff dura mater and insert intracranial electrodes (*10*). To enhance compatibility, we incorporate a stiff shaft (i.e., silica tube) to increase rigidity of the µIP, while maintaining a compliant and micro-invasive diameter (10 µm) 1 – 10 mm at the tip of the probe to avoid significant damage to the recording site. These probes can be applied in standard chambers and with conventional guide tubes, while retaining chronic functionality.

CF is the material of choice for sensors utilizing electrochemical fast-scan cyclic voltammetry (FSCV) as it displays high adsorption, and resulting sensitivity, for dopamine (*11*) . Critically, CF is also capable of recording electrical activity with standard electrophysiology (EPhys) due to its high conductivity (*12, 13*), making it ideal for multi-modal measurements of both chemical and electrical neural activity. FSCV is well established for recording dopamine concentration changes with millisecond resolution and nanomolar sensitivity (*14*) and has been applied across a range of organisms, from flies (*15*) to humans (*8*). This technique involves applying a triangular scan from -0.4 V to 1.3 V at the CF recording electrode tip and measuring reduction and oxidation (redox) current produced by electroactive molecules such as dopamine. Dopamine produces measurable redox currents at potentials of -0.2 and 0.6 V using standard FSCV parameters (e.g., ramp rate of 400 V/s, an Ag/AgCl reference, etc.) (*14*). This recorded current is then used to compute relative fluctuations in dopamine concentration at each time point, using principal component analysis.

FSCV and EPhys recordings interfere with each other, which has traditionally restricted dual use, even when applied at physically separate sensors. This is because FSCV generates voltages that are transmitted through the conductive brain tissue and overshadow signals recorded with EPhys in the form of FSCV artifacts. Methods have recently been developed to overcome this interference by interpolating artifacts in the time and/or frequency domain and enable recordings of both electrical and chemical modes of neural signaling *in vivo* (*7, 16, 17*), which are applied in this work. Techniques also exist to measure these on the same electrode (*18*), albeit with limited sensitivity due to the removal of the negative holding potential between FSCV scans (*19*).

Most *in vivo* dopamine recordings have been applied in rodents and other lower-level species (*6, 15, 20*). Building micro-invasive sensors for nonhuman primates is critical to measure the function of dopamine and other neurochemicals in a species displaying closely related anatomy and behaviors to humans. Enabling these techniques in nonhuman primates will further help strengthen opportunities for clinical translation, such as for providing online measurements of dopamine as biomarkers for controlling therapeutic deep brain stimulation (DBS) in Parkinson’s disease or other disorders known to involve dopamine dysregulation (*21*). A key challenge in translation involves the larger brain size of primates as compared to rodents. Intracranial devices must be long enough to reach deeper brain targets (e.g., the striatum is 10 – 30 mm deep in the rhesus monkey, and 3 – 5 mm in the rodent), while maintaining a sufficiently small diameter to minimize induced implant size-related inflammatory responses, especially surrounding the recording site. Furthermore, the devices should be stiff enough to accurately target regions of interest without significant spatial deflection (*23*).

Dopamine signals in the striatum are known to be spatially heterogeneous, with functional variations observed on a scale as small as tens of micrometers to millimeters across distinct striatal subregions (e.g., dorsal vs ventral or CN vs putamen) (*7, 24, 25*). Such spatial variability in the dopamine signaling emphasizes the need for focal multi-channel measurements. Related to this, the ability to measure neurochemical and electrical signals from more than a single site from a deeply penetrating probe is imperative to identify site-specific signaling patterns and to discriminate heterogenous functions of these signals within a concentrated brain area. The current method to achieve multi-site recording requires multiple individual sensors to be implanted. However, each probe requires an individual penetration, and therefore the total inflicted trauma would scale with each additional implanted sensor. This is likely untenable for achieving high-density recordings safely. Furthermore, standard primate chamber systems require a grid with a discrete inter-electrode spacing, which is usually ≥ 1 mm, which would restrict the ability to record signals from sites within a focal, microscale, region of the brain. Electrode insertions in a standard primate chamber system require larger guide tubes to penetrate the stiff dura mater and safely traverse the delicate CF tip (7 µm diameter). These guide tubes (∼ 0.4 mm diameter) are much more likely to cause hemorrhaging and other traumatic injuries due to their size (*26*). Thus, a single intracranial device that contains multiple recording channels would be advantageous for achieving multi-site measurements within microscale resolution and without significant increases in trauma as used in a chamber or any other acute or chronic cranial window interface.

Recent work demonstrated the use of chamber-compatible s-µIPs in rhesus monkeys (*7*) for chronic neurochemical and electrophysiological recording. These were used to study the relationship between dopamine and beta-band LFPs (*7*) in reward and movement behavioral variables, but the technical fabrication was not detailed in this scientific report. Similarly, the fabrication and application of µIPs developed for rodents have been previously reported (*2, 23*). However, these devices are too short for reaching deep brain targets, such as the striatum, in monkeys and other larger animals, including humans, as explained above, and lengthening such sensors is nontrivial given the flexibility of these sensors, that scales further with length. Here, we detail the fabrication of chamber-compatible s-µIPs as well as the expansion to focal dual-channel recording realized within a single intracranial probe.

## Results and Discussion

s-µIP systems that allow targeting and recording of both neurochemical and electrical neural activity from deep brain regions of primates were successfully created and validated *in vivo* (**Fig. 1**). Single-channel s-µIPs were previously demonstrated for chronic measurements of chemical and electrical neural activity and used to analyze the relationship between striatal dopamine and beta-band LFP signaling in reward and movement variables in monkeys (*7*). Here, we detailed the fabrication methods for these systems, elaborate on *in vivo* performance including demonstrating of spike recording functionality, and expand functionality for dual channel configurations (**Fig. 2**).

**Figure 1.**
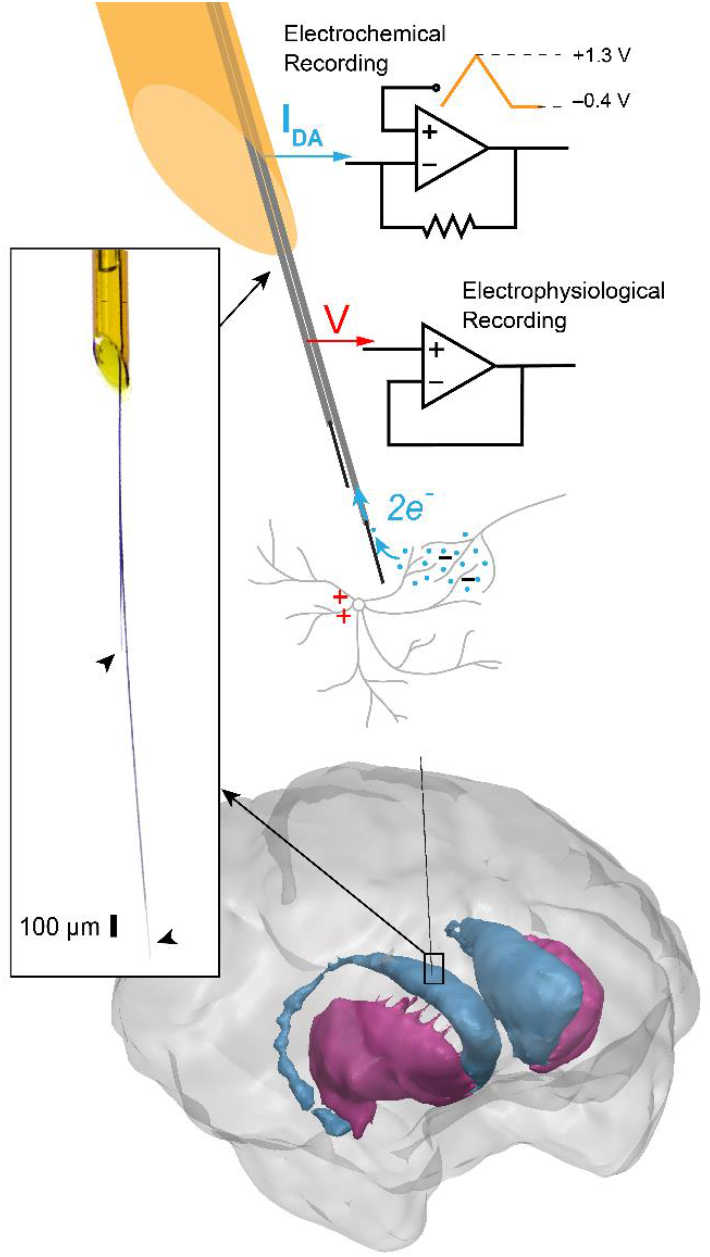
Overview of the s-µIP. The s-µIP is capable of recording both neurochemical and electrical neural activity via electrochemical (FSCV) and electrophysiological (EPhys) recording, respectively. Left inset shows a magnified photo of the tip of a fabricated device, to visualize the recording tips on two flexible CF electrode threads (CFETs) (arrowheads) emerging from the tapered end of the silica tube shaft. The cellular scale (∼ 10 µm) diameter of the CFETs (1 - 15 mm long) ensures minimal trauma to the tissue near the targeted recording site while the stiff shaft (165 µm diameter, 90 - 100 mm long) helps reinforce the device for insertion towards deep brain targets, such as the monkey striatum (illustrated on the bottom).

**Figure 2.**
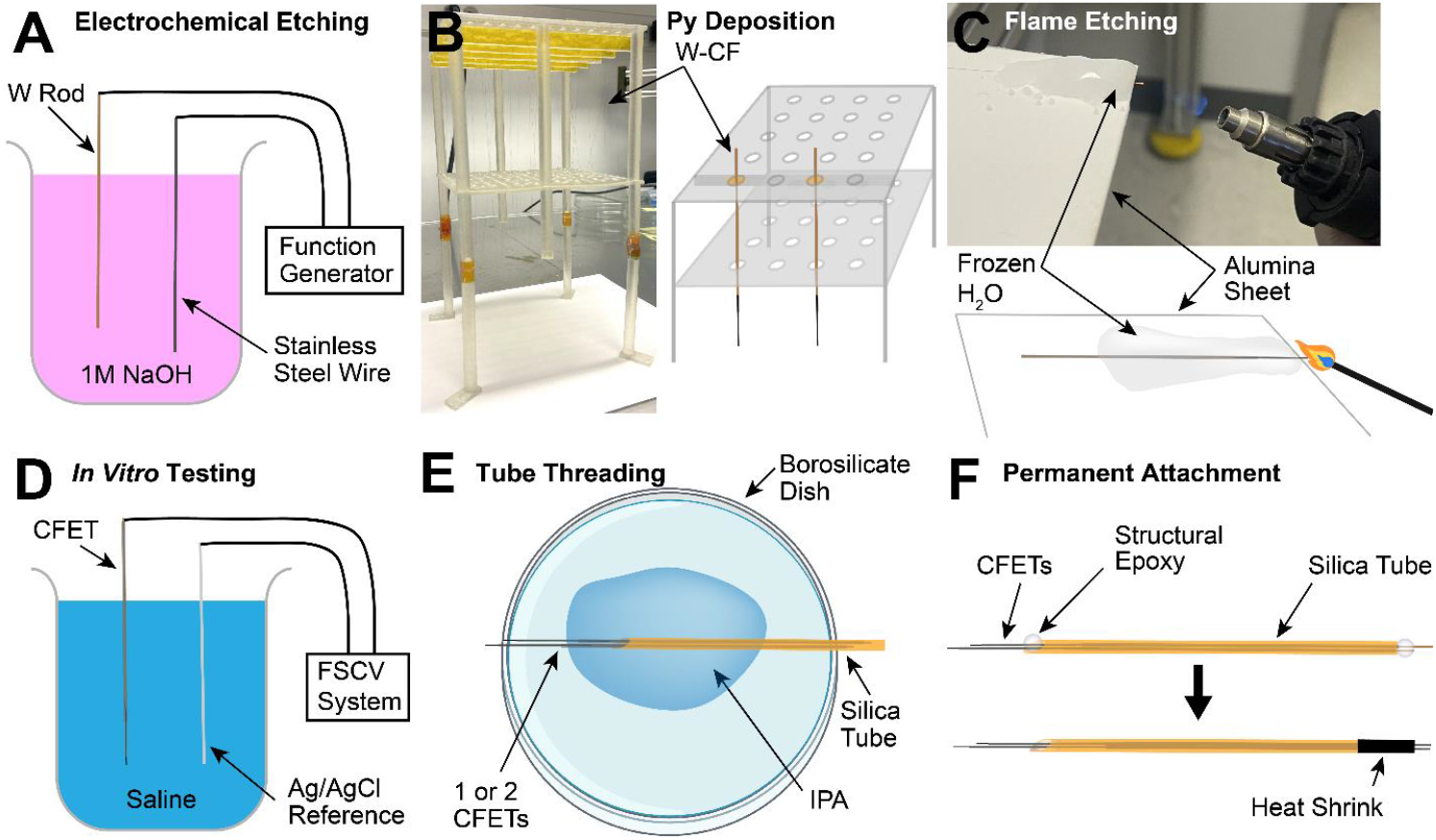
s-µIP fabrication process. Individual steps are further detailed in “Methods: S-µIP Fabrication Process”. (**A**) 150 mm long tungsten (W) rods (50 µm diameter) are electrochemically etched to reduce their thicknesses and create a tapered tip for subsequent connection of the carbon fiber (CF). (**B**) The tungsten wire is attached to a 20 – 30 mm long CF using silver epoxy. Multiple tungsten-CF assemblies (W-CF) are mounted on a custom 3D-printed filament separating fixture and are processed for conformal parylene-C (Ppy) deposition. (**C**) The py insulation is stripped from the tips of the coated CFs (py-CFs) to expose the recording tips by flame etching with a torch while maintaining thermal insulation of the remainder of the probes using a freeze-anchoring platform. (**D**) The assembled and patterned CF electrode thread (CFET) is tested *in vitro* in 0.9% saline to validate dopamine detection functionality. (**E**) Single or pairs of CFETs are threaded into silica tubes (165 µm outer diameter) with isopropanol (IPA) to reduce friction and facilitate the threading process. (**F**) The CFETs are permanently attached to the silica tube using structural epoxy. The tail end of the CFETs are stripped using flame-etching and used to connect to external instrumentation (not shown). These exposed wires are further protected with heat shrink tubing.

A key change required for fabrication of the s-µIPs, as compared to other similar micro-invasive sensors used for smaller animals (i.e., rodents) (*2, 20, 23, 27*), was the significant elongation of the device to target deeper brain structures in larger animals (i.e., monkeys). Thus, the fabrication process was redesigned to allow for practical manufacturing and handling of such elongated CFs. This included creating a filament separator fixture to prevent CFs from attaching to each other due to electrostatic interactions and/or Van der Waals forces, and freeze-anchoring techniques to stabilize the significant tendency of the thin CFs to displace due to surface tension and capillary effects at the air-water interface during flame-etching (**Fig. 2**). These methods were not required for shorter µIP implants used for rodents (*20*) where striatal targets lie more superficially (i.e., 3 – 5 mm deep), but were needed to reach homologous regions that are much deeper (i.e., > 15 – 30 mm deep) in the monkey.

We aimed to keep the diameter of the sensors the same as what was used in the rodents (i.e., 10 µm), and as small as possible to minimize induced trauma in the brain tissue. At the same time, stiffening of a large portion of the implant was necessary to allow for successful insertion to the deep striatal targets in the monkey, using standard primate recording chambers and guide tube insertion protocols to cross over the stiff outer meningeal membranes. Thus, we designed the length of the flexible parylene-coated CF (py-CF) (i.e., the “micro-invasive” portion of the device) to range between 5 – 15 mm. The remaining length (∼ 100 mm) of the device was ensheathed with a larger silica tube (165 µm diameter) shaft to increase stiffness for manual handling and brain insertion. Creating a discrete length of py-CF was imperative to isolate the recording site (i.e., the tip of the py-CF) from the larger and stiffer shaft. Devices with a similar diameter and stiffness to our shaft are known to produce tissue responses that spread several hundreds of microns (*2, 28*). Thus, separation from the stiff shaft was designed to restrict spreading of the shaft’s induced tissue responses to the recording site and ensuring functional recording of neural signals in a minimally obstructed volume of tissue. However, even with the addition of the stiff shaft, insertion yields remained low ranging from 21.4% (3/14 individual sensors) to 33% (3/9). The primary mode of failure was the py-CF fracturing due to lateral bending forces that increase the probability of buckling as the probe is advanced through the guide tube and into brain tissue. These effects were found to be exacerbated with longer py-CF lengths. Reducing the length of the py-CF may help to reduce these failures, and examining the functional performance as a function of this length may help optimize these design parameters in future work. Also, automating the insertion process by using a motorized micromanipulator, rather than manual lowering procedures, to control the insertion speed may help reduce the risk of buckling. Initial experiments in brain phantom (i.e., 0.6% agar) demonstrate that slower insertion rates (≤ 0.2 mm / s) diminished electrode buckling.

*In vivo* device performance of the s-µIPs was validated by measuring reward- and movement-related changes in dopamine, LFP, and spike activity in two monkeys. Devices were implanted into the striatum (caudate nucleus, CN, or putamen) and recordings began 2 – 3 months post-implant. Monkeys were trained to perform tasks to measure reward and movement related dopamine signaling and corroborate findings with prior work. These eye movement tasks involved making saccades and fixating on a series of two cues (i.e., central cue followed by a peripheral value cue) displayed on a screen in front of them to obtain rewards in the form of liquid-food (monkey P) or diluted apple juice (monkey T). In the “direction task”, only one value cue was displayed on each trial and the direction (left or right, or top or bottom) of the cue indicated the reward size (large or small). In the decision-making “shape task”, one (forced trials) or two (choice trials) cues were displayed on each trial and the shape of the cue represented the upcoming reward size (large or small). Example measurements from single trials in the decision-making shape task demonstrate clear time-varying changes in dopamine, as recorded in the CN (**Fig. 3A**). These fluctuations are apparent especially after value cue presentation, where we would expect to see reward associated dopamine changes, given dopamine’s well established role in reward prediction error and motivation (*29*). These measurements also demonstrate the ability to measure dopamine from two neighboring channels on a single s-µIP with 0.1 – 0.2 mm separation between the recording tips. Single-trial LFPs measured from another s-µIP are shown to display fluctuations in the frequency of beta-band “bursts” (i.e., 1 – few cycles of high beta-band power) around the cue and other task events, as expected based on previous reports (*30*) (**Fig. 3B**).

**Figure 3.**
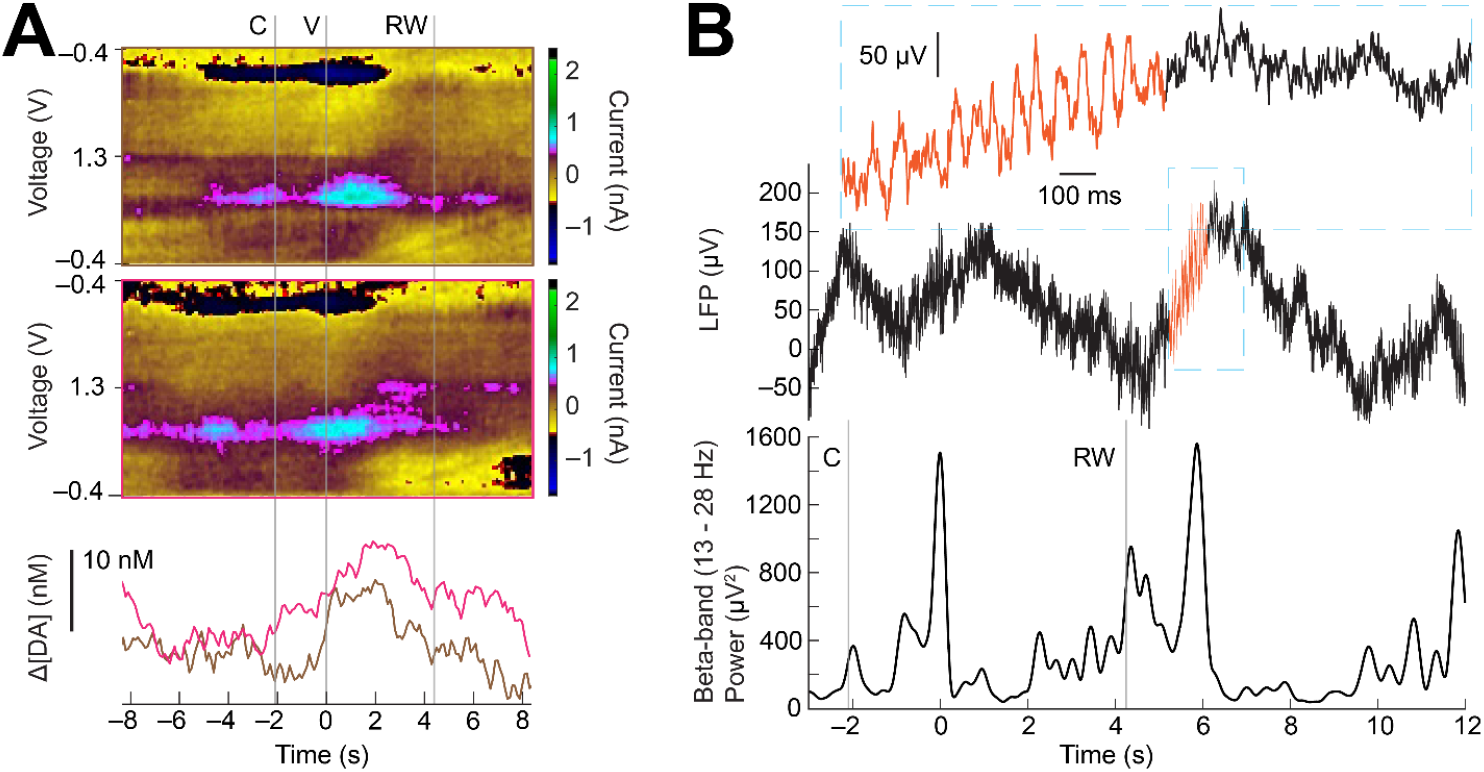
*In vivo* FSCV and EPhys measurements from s-µIP from task-performing monkey T. (**A**) FSCV data collected from two neighboring sites using a dual-channel s-µIP, showing single-trial measurements as the monkey performed a forced choice large reward trial in the shape task (sites c8dg and c8ds: 483 days post-implant). Color plot shows clear dopamine redox current (i.e., color changes ∼ 0.6 and -0.2 V). PCA extracted dopamine concentration change ([ΔDA]) is plotted below the color plot and highlights changes around task events (e.g., increases in both channels after the value cue, V). (**B**) Single-trial LFP signals measured as monkey performed a large reward trial (top panel) in the direction task, with time plotted relative to value cue display at 0 s (site c3bs-c3a: 496 days post-implant). Top inset displays a close-up of the LFP signal where beta bursts are visible (orange traces). The bottom panel is the beta-band power to highlight task relevant changes in beta signaling. For all figure panels, C indicates central cue, V is value cue display, and RW is reward outcome.

Data were averaged across trials to distinguish patterns of neural signaling between distinct reward (large vs. small) and movement (contralateral vs. ipsilateral cues) conditions during the value cue fixation window, where monkeys could anticipate the forthcoming reward based on the value cue identity (i.e., shape) and made an eye movement in a distinct direction. For example, an s-µIP in the CN displayed increased levels of cue-evoked dopamine for large reward conditions as compared to small reward trials (**Fig. 4A**), as would be expected based on the canonical role of dopamine in reward. Beta-band LFP, as measured from another s-µIP, also displayed reward-related modulation, albeit in the opposite direction, showing enhanced suppression (i.e., event-related desynchronization, ERD) in response to the cue (**Fig. 4B**). This relationship was expected based on prior studies showing that beta-band signaling shows reward modulation in the opposite polarity (i.e., more suppression for larger reward value) (*31, 32*). This inverse relationship between reward related dopamine and beta-band LFP has been previously characterized in similar tasks using standard CFE sensors (*32*). Beta-band LFP showed enhanced suppression when saccades were made to contralateral cues relative to ipsilateral cues (**Fig. 4C**). The lateralized signaling for striatal beta-band LFP, as well as dopamine (not shown here), has also been shown in previous work (*32*). Spike measurements from an s-µIP in the CN of monkey P also displayed reward-modulated differences in firing rate during the value cue fixation window, similar to the dopamine measurements, and also to previous findings of CN activity related to reward (*16, 31*) (**Fig. 5**).

**Figure 4.**
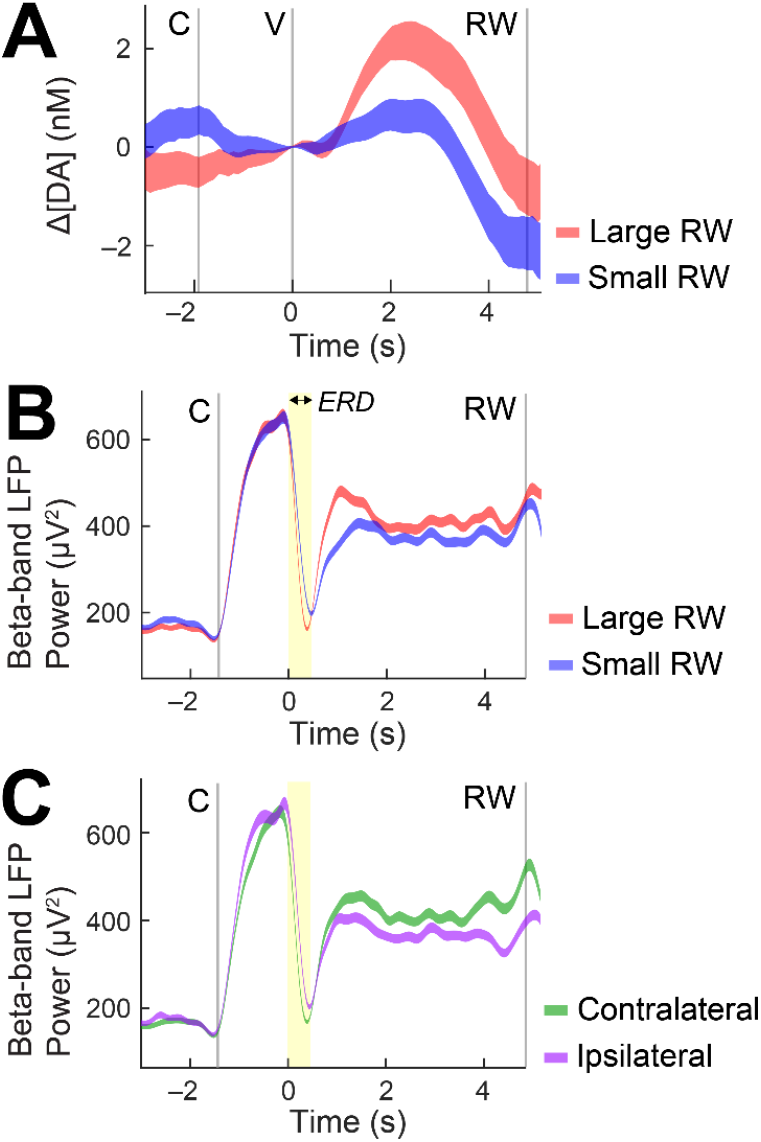
Dopamine and LFP activity related to specific behavioral conditions in monkey T. (**A**) Trial-averaged dopamine for large (red) and small reward (blue) conditions for successful forced choice trials in the shape task. Higher dopamine levels are observed following the display of large reward associated value cues as compared to small reward cues in monkey T (site c8dg: 483 days post-implant). (**B**) Trial-averaged beta-band LFP power for large and small reward conditions in the direction task, showing increased suppression (i.e., ERD, highlighted in yellow) for large as compared to small reward cues during the brief time window following the value cue at 0 s (site c3bs-c3a: 496 days post-implant). (**C**) Trial-averaged beta-band LFP power for a fixed large reward condition for left (contralateral) and right (ipsilateral) cues for the same session and site as (B). Increased suppression for contralateral is observed as compared to ipsilateral movement conditions. For all figure panels, C indicates central cue, V is value cue display, and RW is reward outcome.

**Figure 5.**
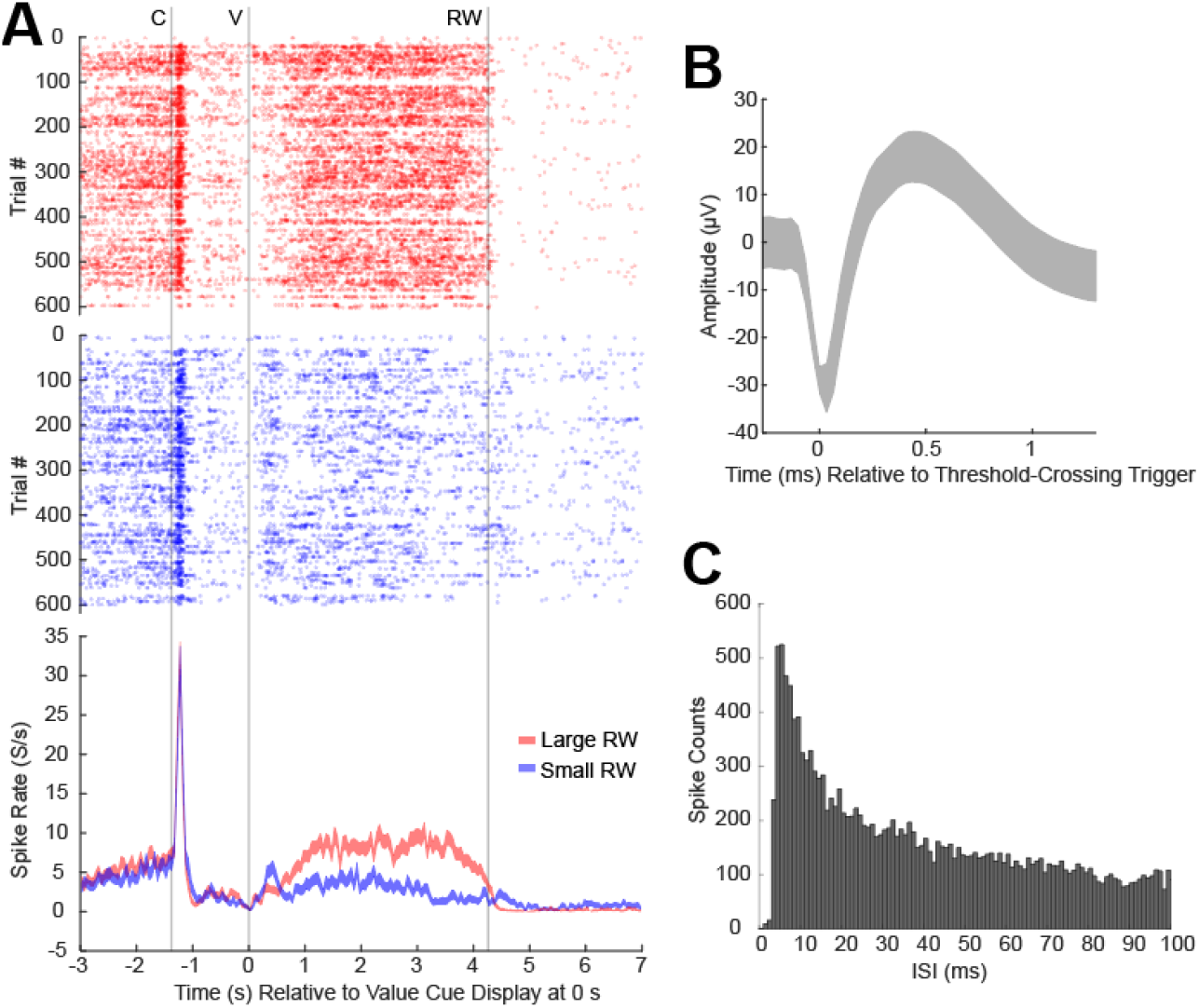
Spike activity recordings from s-µIP implanted in monkey P in the direction task (site c33: 390 days post-implant). (**A**) Raster plot showing detected spikes as a function of time relative to the value cue display (V at 0 s) for each trial (y-axis) with peristimulus time histogram (PSTH) plotted in the bottom panel showing average spike firing rate (y-axis) for the large (red) and small (blue) reward conditions relative to the same V event. C is the central cue and RW is reward outcome. (**B**) Average spike waveform +/-standard deviation. (**C**) Interspike interval (ISI) histogram showing the distribution of the timing between spikes detected. Bin widths are 1 ms.

Measurements were made from chronically implanted s-µIPs that maintained functionality > 1 year post-implant. Preliminary measurements demonstrated that these sensors maintain longitudinal performance in dopamine detection (*7*), but more work is needed to characterize functional operation over a larger number of devices and longer time periods, and as a function of FSCV recording durations (e.g., cycles of 10 Hz triangular waveforms applied). Next steps will include enabling multi-modal measurements from a single probe and characterizing the magnitude of interference between EPhys and FSCV recording channels as a function of channel separation on the s-µIP. Future work will also entail improving the insertion yield and increasing the channel capacity of the s-µIP to enhance our ability to capture the heterogeneous signaling patterns displayed in focal neural circuits.

## Methods

### s-µIP Fabrication Process

The fabrication of the s-µIP involves manipulation of microscale materials in a clean environment under a stereomicroscope as well as electrochemical etching processes that are performed in a fume hood. The first step involves forming the conductive wire that will provide an electrical connection between the CF sensing material and the external FSCV and EPhys instrumentation. Typical microelectrodes used in primate electrophysiology usually range between 80 – 150 mm in length, despite the targeted brain depth usually being < 50 mm. This additional length is necessary, in some cases, to attach the electrodes to an external micromanipulator that is anchored to the implanted chamber so that they can be carefully lowered into the chamber grid. The grid is a plate with discrete holes that allows the precise mapping of electrode coordinates from MRI images of the brain and the insertion of the electrodes into the exposed brain. In other cases, this additional length is necessary to attach the electrodes to microdrives installed on the chamber grid and connect the electrodes to electrode interface boards mounted on the chamber (*33*). A tungsten (W) rod (50 µm in diameter and 150 mm in length) (A-M Systems, 715550) is chosen due to its high tensile strength and Young’s modulus (E = 3 – 4 × 10^11^ N / m^2^), ensuring a robust and stiff shaft. The W rod is etched down to a 20 – 30 µm diameter with a slight taper (over a length of 2 – 3 cm) on the front end (where the CF will be attached) to prevent additional thickening due to CF attachment. Electrochemical etching is performed in 1 M sodium hydroxide (NaOH) by applying a 15 V_pp_ 1 kHz sinusoidal waveform onto the wire (**Fig. 2A**). A “dip coater” system (Nilo Scientific, Ni-Lo X2 Dip Coater) is used to control the etch rate as a function of length by elevating the wire slowly out of the solution during the etching process.

A 20 – 30 mm long CF (7 µm diameter) (Goodfellow, Grade 34-700, CAS# 7440-44-0) is attached to the tapered end of the W using silver epoxy (Epo-Tek, H20S) that has been diluted with isopropanol. The CF overlaps with the W wire 10 – 15 mm to provide a sufficiently long conductive path between the two elements. The assembled CF-W electrode is placed in an oven at 80°C for 3 hours to cure the silver epoxy. The CF-W is then coated conformally with 0.5 – 1.3 µm of parylene-C to provide electrical insulation (Specialty Coating Systems, SCS PDS 2010) (*23, 32*). Additional procedures and materials are required to manipulate and process these longer and more flexible electrodes as compared to prior devices used for smaller animals (*20, 23*). 20 – 30 CF-W assemblies are arranged onto a custom-made filament-separating fixture to prevent the CF-W’s from attaching to each other during transportation and parylene deposition (**Fig. 2B**). The surface of the CFs inherently adsorb to other surfaces and to themselves due to electrostatic, Van der Waals, and/or other external forces (such as ventilation or air handling), and these effects become more pronounced with longer probes.

The parylene-coated CF-W’s are then etched to expose a 100 – 300 µm length of bare CF to serve as the recording tip (**Fig. 2C**). This functional assembly is termed CF electrode thread (CFET), to be consistent with prior work (*23*) and to distinguish it from the final s-µIP device that includes the protective silica sheath. A freeze-anchoring technique is used to thermally insulate the CFET as well as anchor it in place while a butane blow torch is used to thermally etch the exposed tip. This involves pouring distilled water onto a thermally stable ceramic plate (e.g., alumina) with the CFET, and then freezing the water with dry ice. A few millimeters of the parylene-coated CF (py-CF) portion of the device remains exposed, protruding out of the ice and off the edge of the plate. The flame of the blow torch is then directed towards the exposed py-CF to thermally decompose the parylene-C and expose a discrete length of bare CF. The ice is allowed to thaw and then the etched CF may be further trimmed to obtain optimal current if needed after subsequent *in vitro* measurements.

The tail-end of the etched CFET (i.e., tungsten wire), is also etched with the blow torch to remove the overlying parylene insulation and expose the bare W wire. The exposed W is then crimped to a Mill-Max pin connector (Mill-Max, 0489-0-15-01-11-02-04-0) and connected to an FSCV current-to-voltage transducer for electrochemical measurements., Then, the tip of the CFET is submerged in 0.9% saline to record FSCV current, allowing us to record the background current amplitude and noise levels, which are indicators of the sensitivity and limit of detection of the sensor (**Fig. 2D**) (*20, 34*).

One or two functional CFETs are then threaded through fused silica tubes (100 – 165 µm outer diameter) cut to 9 – 10 cm lengths (Molex, Polymicro, TSP100170) (**Fig. 2E**). The front-end of the silica tube is cut with a taper (30 – 45 degrees bevel) using a dicing machine to facilitate subsequent brain insertion. The crimped connector is removed from tested CFETs during the threading process as this tail-end is threaded first into the silica tube. Isopropanol is used to reduce the friction between the CFET and the silica during the threading process. 0.2 – 5 mm of the py-CF is concealed in the tube, and 1 – 15 mm of the py-CF is exposed, protruding outside of the tube. A longer length of free-hanging py-CF is advantageous in reducing size-related implant-induced trauma, but at the expense of increasing mechanical failure due to buckling during brain insertion. This compromise was discussed further in the “Results and Discussion” section. Threading of multiple CFETs occurs at the same time in the tube, since it is more difficult to insert a second electrode when the tube is already threaded with an electrode. Separation between the two CFET tips was manually provided by pulling the tail ends of the electrodes until the targeted separation was made. This separation ranges from 0.1 µm - several millimeters. The electrodes are permanently attached to the tube using structural epoxy (Devcon, 14250) on both ends of the silica tube (**Fig. 2F**). Finally, polyolefin heat shrink tubing (Raychem, Microfit, 0.365 mm expanded diameter, 0.178 mm recovered, MFT-#1×4’-BLK) was used to reinforce the tail-end of the CFETs coming out of the silica tube. Final assembled s-µIPs are tested again *in vitro* to ensure functional performance.

### Electrode Implant Procedures

Two rhesus monkeys were used for *in vivo* validation of the s-µIPs described in this work. Each monkey underwent 3 separate procedures for installation of the cranial chamber, a craniotomy to expose the brain within the chamber, and chronic electrode implantation through the chamber. These are described in detail elsewhere (*32*) and protocols are also published online (dx.doi.org/10.17504/protocols.io.kqdg32b91v25/v1, dx.doi.org/10.17504/protocols.io.x54v92wd4l3e/v1, dx.doi.org/10.17504/protocols.io.bp2l62m95gqe/v1). All animal procedures were approved by the Institute’s Animal Care and Use Committee (IACUC) at the University of Pittsburgh, by the Committee on Animal Care (CAC) of the Massachusetts Institute of Technology, and performed following the Guide for the Care and Use of Laboratory Animals (Department of Health and Human Services), the provisions of the Animals Welfare Act (USDA) and all applicable federal and state laws in Pennsylvania and Massachusetts. Here, we summarize the procedure for implanting the s-µIPs into the chamber and to targeted brain areas in the striatum.

The electrode implantation procedure involved implanting both s-µIPs and standard CF electrodes (CFEs). Electrodes were threaded through guide tubes (Connecticut Hypodermics, 27G XTW “A” Bevel) that were pre-filled with sterile petrolatum lubricant (Dechra, Puralube Vet Ointment) and attached to a microdrive. Electrodes (e.g., s-µIPs) were lowered into targeted holes of a grid installed on top of the chamber. The relative positions between the grid holes and the brain were estimated based on co-registered images using MRI and CT. This allowed us to insert sensors to predetermined brain areas in the CN and putamen. During insertion, this microdrive-electrode assembly was positioned over the targeted grid hole and manually lowered onto the grid. The petrolatum lubricant helped to keep the electrode and guide tube attached to each other during lowering through capillary forces. This lubricant also served to reduce friction when the electrode is lowered through the guide tube and prevent fluid from inside the brain from channeling up the guide tube. During the initial microdrive lowering, the s-µIPs remained fully retracted inside the guide tubes to protect the delicate CF tip from breaking when penetrating the stiff dura mater. The guide tubes first pierced the dura mater and were lowered into the brain until they were 5 – 10 mm above the targeted recording sites. This depth was achieved by cutting each guide tube to a predetermined length from the targeted depth to 2 mm above the grid. The microdrive was then secured to the grid with acrylic cement and all guide tubes were elevated until they were just above the brain surface.

### FSCV and EPhys recordings

During each recording session, a 4 channel FSCV system was used to record dopamine from a subset of implanted electrodes, and an EPhys system (Neuralynx, Digital Lynx SX) was connected to all remaining electrodes, as done in previously reported experiments (*16, 32*). The Ag/AgCl electrodes implanted above the epidural tissue and inside the EPhys were connected to the FSCV system ground and used as the reference for both the FSCV and EPhys recordings.

FSCV operates by applying a triangle waveform from –0.4 V to 1.3 V at a rate of 400 V/s and measuring current generated by reduction and oxidation (redox) of molecules on the surface, including dopamine, which displays redox at –0.2 and 0.6 V. The triangular waveform is applied every 100 ms for an effective sampling frequency of 10Hz. The recorded voltage-dependent current or cyclic voltammogram (CV), is then concatenated at each 100 ms sampling interval to create a 2-dimensional plot where the current is plotted as color, voltage is represented on the y-axis, and time on the x-axis. Such color plots are useful to evaluate the voltage-dependent current changes and confirm selective dopamine redox. Recorded electrochemical current was background-subtracted at relevant timepoints (e.g., value cue) to analyze neural signaling changes relative to specific behavioral states, as described in “Analysis” below. This subtraction is necessary as FSCV with standard CF-based sensors inherently generates current drift over periods > 30 – 60 s, which can overshadow behaviorally relevant signals.

EPhys recordings were made using a unity-gain analog amplifier headstage (Neuralynx, HS-36) with an input range of ±1 mV at a sampling frequency of 30 (monkey P) or 32 kHz (monkey T), and bandpass filtered from 0.1 Hz to 7500 Hz. This system also recorded timestamps of identified task events using 8-bit event codes. The EPhys and FSCV systems were synchronized by transmitting uniform “trial-start” event codes to both systems, as detailed in previous work (*16, 32*).

### Behavioral Tasks

FSCV and EPhys recordings were made from the striatum as monkeys performed a simple reward-biased eye movement task, that we refer to as the “direction task” for simplicity here, and as described previously (*7, 35*). Each trial of the task began with a central fixation cue being presented to the monkey on a screen. After a two second fixation, a value cue was presented either on the top or bottom (both left of center, contralateral to implanted electrodes) of the screen. In some sessions, the value cue was presented on the left (contralateral) or right (ipsilateral) of the screen. The location where the cue appeared on indicated whether a big or small reward would follow. The big and small reward locations were changed on every block. The monkey was trained to saccade to the value cue and hold fixation for 4 seconds, after which she received the reward associated with that value cue.

Recordings were also made, in monkey T, in a simple variation of this task that incorporated more explicit decision-making variables. In this second task variation, referred to here as the “shape task”, two value cues were presented on each trial, with each cue having a distinct shape. The shape of the cue was associated with the reward size (large or small) and animals learn to identify and choose the shape that offers the larger reward through trial and error. Forced choice trials were also interleaved (25% of all trials) where only one value cue was displayed, and the monkey was forced to choose the displayed large or small value cue. The cue-reward association was reversed after a block of 50 – 300 trials depending on the session. Only forced choice trials are shown in results plotted herein.

### Analysis

All FSCV and EPhys signals were analyzed in MATLAB (MathWorks, MATLAB 2021b) as previously described (*16, 32*).

Dopamine concentration changes (Δ[DA]) were extracted from the background-subtracted FSCV recorded currents using principal component analysis (PCA) (*3*). Task modulated Δ[DA] related to the value cue was computed from the background-subtracted current. The background-subtracted current was derived by subtracting the recorded current by the current at the value cue display event, which is the start of the targeted timeframe of interest, after which, we expect reward or movement related signals. All Δ[DA] signals plotted herein are computed with this same reference point for background subtraction. This background-subtracted current was projected onto the principal components computed from standards of dopamine, pH, and movement artifacts (*3, 7*). This then allowed us to estimate the current contributions associated with dopamine, pH, and movement. Signals that displayed a variance above a threshold (i.e. Q > Q_α_ where Q is the residual sum of squares and Q_α_ is a calculated threshold (*36*)) are automatically assigned null values (i.e. NaN in MATLAB) and not used in resulting calculations. This reduces the contribution of extraneous signals such as charging artifacts due to movement or other sources of noise (*3*). Additionally, Δ[DA] signals were nulled wherever CVs showed a correlation greater than 0.8 to movement standards (*3, 7*) to reduce the possibility of falsely attributed dopamine signals.

Beta-band LFP was computed from bipolar EPhys measurements made by pairs of electrodes during concurrent FSCV recording, as described previously (*7*). FSCV artifacts were removed from the EPhys measurements using spectral interpolation and then downsampled to 1 kHz. Signals were bandpass filtered around 13 to 28 Hz, squared, enveloped, and then smoothed with a Hanning window of 0.25 s, as done previously (*7*).

Spike activity was analyzed from the same EPhys data, but were referenced to the original ground reference rather than another electrode pair since local referencing is more critical for localizing the source of LFP signals. FSCV artifacts were removed using time-domain interpolation algorithms (*16*) as to preserve the high frequency content of the signal for standard spike sorting. Spike sorting was performed in software (Plexon, Offline Sorter), following previously described techniques (*16*).

## Data Availability

Protocols used in this study are available at protocols.io

(dx.doi.org/10.17504/protocols.io.kqdg32b91v25/v1,

dx.doi.org/10.17504/protocols.io.x54v92wd4l3e/v1,

dx.doi.org/10.17504/protocols.io.bp2l62m95gqe/v1).

## Code Availability

MATLAB code used to analyze data may be found at GitHub as made available through zenodo (https://doi.org/10.5281/zenodo.10955583, https://doi.org/10.5281/zenodo.10397773).

## Author Information

3501 Fifth Avenue; 4071 Biomedical Science Tower 3; Pittsburgh, PA 15213.

## Author contributions

H.N.S., U.A., J.C., S.M., R.S., R.M., A.S., and O.C. designed and performed research. H.N.S., U.A., and S.M. analyzed data. H.N.S., U.A., and S.M. wrote the paper.

## Acknowledgment

The authors thank Ms. Rebecca Marflak, Dr. Esta Abelev, Dr. Daniel Lamont, Dr. Julia K. Oluoch, Ms. Erica N. Griffith, Ms. Baylie S. Leveto, and animal care staff at the Systems Neuroscience Center Animal Resource Laboratory (SNARL) at the University of Pittsburgh (Pitt). Work performed at Pitt’s Nanofabrication and Characterization Core Facility (RRID:SCR_05124) and services and instruments used in this project were graciously supported, in part, by Pitt.

